# Targeted Bacteriophage T4 Nanoparticles Reverse HIV-1 Latency in CD4+ Human T Cells

**DOI:** 10.1101/2021.07.20.453091

**Authors:** Himanshu Batra, Jingen Zhu, Swati Jain, Neeti Ananthaswamy, Marthandan Mahalingam, Pan Tao, Camille Lange, Shawn Hill, Chaojie Zhong, Mary F. Kearney, Haitao Hu, Frank Maldarelli, Venigalla B. Rao

## Abstract

A major barrier for HIV-1 eradication is the latent virus reservoir containing stably integrated and silent proviruses in CD4+ T-cells. Targeted reactivation and removal of this latent reservoir is a potential strategy for HIV-1 cure but remains a major challenge. Here, we investigated whether CD4-targeted bacteriophage T4 capsid nanoparticles that mimic HIV envelope can reactivate HIV-1 latency. The nanoparticles were arrayed with CD4-binding CD4-DARPin, or HIV-1 gp140 envelope trimer. When exposed to J-Lat T-cell model of HIV-1 latency or primary T-lymphocytes from human PBMCs, these nanoparticles activated CD4+ T-cells without causing global T-cell activation, which led to activation of HIV-1 proviral transcription, viral protein production and release. Intriguingly, the observed T-cell activation and HIV-1 latency reversal do not involve the classic PKC or NFAT pathways and did not lead to cytokine storm. These studies indicate that engineered non-infectious bacteriophages can be exploited for HIV-1 cure and targeted T-cell therapies.

## Introduction

About 40 million people worldwide are currently living with HIV-1, and 1-2 million new infections occur every year(’UNAIDS (2024). UNAIDS DATA 2023 Accessed 10/02/2024’). At least two-thirds of them have access to antiretroviral therapy. ART is lifesaving, though it does not cure HIV-1 infection and is associated with long-term adverse effects(O’Brien et al. 2003).New strategies to eradicate or suppress HIV-1 without ART are, therefore, a global health imperative.

During HIV-1 infection, reverse-transcribed viral DNA integrates into the host genome (provirus) and persists in cells for the lifetime of the cell. Most infected CD4+ T-cells actively produce new viruses and are rapidly eliminated, but in a fraction of infected cells, the proviruses become quiescent. These cells may undergo clonal expansion, but HIV remains largely transcriptionally silent(Churchill et al. 2016; Maldarelli et al. 2014; Musick et al. 2019). Viral rebound rapidly occurs after a pool of proviruses reactivate when ART is discontinued(Chun et al. 1999). Thus, reducing or eliminating this latent reservoir is essential to achieve “remission” or “cure”.

One of the potential curative strategies is “shock-and-kill”, which involves reversing the latent state by inducing HIV-1 transcription with subsequent elimination by cytopathology or by the immune response(Deeks 2012). Although several shock-and-kill agents can reactivate HIV *in vitro*, clinical studies have not demonstrated broad HIV reactivation *in vivo*, or reduction in replication-competent reservoirs(Sloane et al. 2020). The reasons for this lack of efficacy are uncertain and remain as the most difficult obstacle. New approaches, particularly those that can also precisely target therapy to CD4+ T cells are of great interest because these might potently reactivate HIV-1 from latency and eventually lead to removal of the proviral reservoir.

We took an Innovative approach to address this question by engineering a bacteriophage (phage), namely T4, to mimic the envelope of HIV-1 virion and target it to CD4+ T cells to reactivate the latent provirus. Phage T4 belongs to the *Straboviridae* family and infects the *Escherichia coli* bacterium(Walker et al. 2022). T4 is endowed with certain unique features for its engineering as a targeted nanoparticle to CD4+ T cells(Miller et al. 2003). It has a large 120 x 86 nm prolate icosahedral capsid (head) into which 1̃71 kb linear dsDNA viral genome is packaged, powered by a molecular machine assembled at the portal vertex(Chen et al. 2017; Fokine et al. 2004) (Figure 1A). Its outer surface is arrayed with two non-essential capsid proteins; 870 molecules of 9.1-kDa Soc (small outer capsid protein) and 155 molecules of 40.4-kDa Hoc (highly antigenic outer capsid protein)(Fokine et al. 2004; Qin et al. 2010; Yanagida, Suzuki, and Toda 1984). While Soc assembles as trimers at the quasi-three-fold axes and forms a “cage” around the capsid reinforcing the structure, Hoc is a 180Å-long monomeric fiber extending from the center of each capsomer(Fokine et al. 2004; Fokine et al. 2023) (Figure 1B). We engineered Hoc and Soc to generate HIV-mimicking T4 nanoparticles (NPs). We also reasoned that CD4 targeting molecules attached to the N-terminal tips of the 18nm flexible Hoc fibers (shown in Pink, Figure 1B) will have considerable reach and avidity, that can help navigate the T4-NPs to CD4+ T cells.

**Figure 1.**
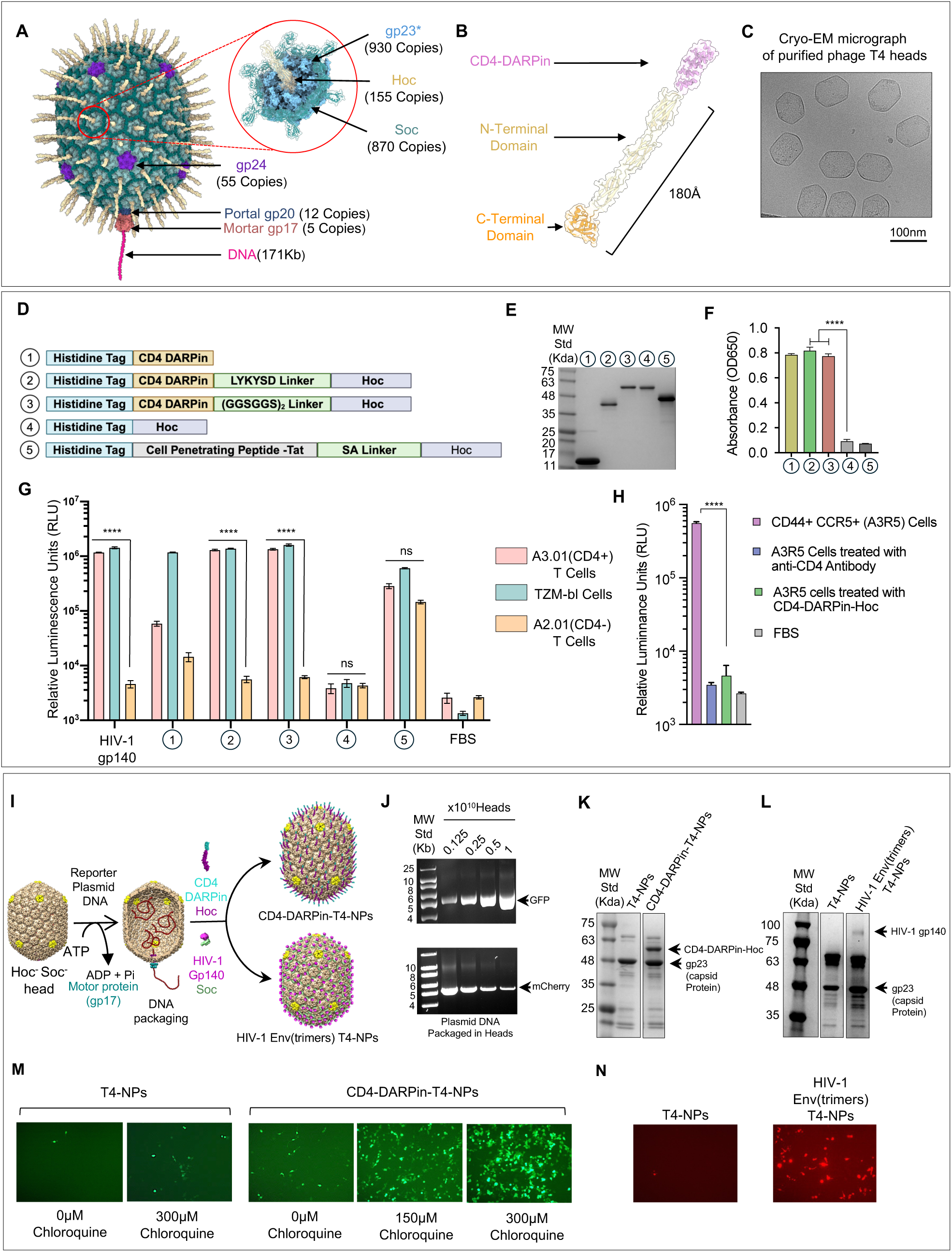
Displaying CD4-DARPin or HIV-1 Env trimers on T4 capsids targets the nanoparticles to CD4+ T cells. (A) Structural model of phage T4 head (capsid). One enlarged capsomer (hexamer) shows the arrangement of the major capsid protein gp23* (Blue), vertex protein gp24* (purple), Soc trimers (Teal), Hoc fiber (Yellow), portal gp20 (Dark blue), and pentameric DNA packaging motor protein, gp17^26^ and DNA (Magenta). (B) Structural model of CD4-DARPin-Hoc fusion protein depicting CD4-DARPin fused to the N-terminal tip of 180Å-long Hoc fiber. (C) Cryo-electron micrograph of purified T4 heads. (D) Schematic of CD4-DARPin-Hoc and HIV-1 gp140-Soc fusion constructs. The DARPin coding sequence was fused to the N-terminus of Hoc via linkers. The unfused CD4-DARPin, Hoc, and CPP-T-Hoc are controls. A hexa-histidine tag (blue) was fused to the N-terminus of all proteins. (E) SDS-PAGE showing purified recombinant fusion proteins. (F) ELISA assays showing the targeting specificity of CD4-DARPin fused proteins to human CD4 receptor. (G) Specificity of CD4-DARPin-Hoc binding to CD4 receptor on T cells. Binding of CD4-DARPin-Hoc variants (panel D) and HIV-1 gp140 Env trimers to CD4+ (A3.01), CD4- (A2.01), and CD4 overexpressing (TZM-bl) cells were quantified. (H) CD4-DARPin-Hoc fusion protein blocked the binding of HIV-1 Env trimers to A3R5 CD4+ T cells in cell binding assay. (I) Schematic representation of *in vitro* packaging and display of CD4-DARPin-Hoc and gp140-Soc timers on T4-NPs. (J) *In vitro* DNA packaging of Hoc-Soc-phage T4 heads with linearized 5.4kb eGFP reporter plasmid. (K) SDS-PAGE showing *in vitro* display of CD4-DARPin-Hoc protein on T4 capsids. The arrows show the band positions of capsid-bound CD4-DARPin-Hoc and gp23 capsid protein. (L) *In vitro* display of HIV-1 gp140-Soc trimers. The arrows show the band positions of capsid-bound HIV-1 gp140-Soc and gp23. (M) eGFP plasmid-packaged capsids decorated with or without CD4-DARPin-Hoc were used for targeted delivery. Fluorescence micrographs of CD4+ 293T cells showing delivery and expression of eGFP transgene without chloroquine and with chloroquine at different concentrations as shown. (N) mCherry plasmid-packaged capsids decorated with or without HIV-1 gp140-Soc were used for targeted delivery. Fluorescence micrographs of CD4+ 293T cells showing the expression of delivered mCherry plasmid. Error bars in various panels show S.D. The P-values were determined using a two-tailed, unpaired t-test. **** = P value <0.0001. Error bars show the S.D. P-value was determined using a two-way ANOVA test. **** = P value <0.0001.

Here, we report proof-of-concept data for the design of human CD4 receptor-targeted T4-NPs and their ability to reactivate the HIV-1 provirus present in the latent CD4+ T cells. We created such targeting particles by decorating the T4 capsid with a high affinity CD4-binding DARPin (Designed Ankyrin Repeat Protein) fused to Hoc, or with a native-like HIV-1 envelope glycoprotein (Env), the SOSIP gp140 ectodomain trimer fused to Soc(AlSalmi et al. 2015; Ringe et al. 2013). Remarkably, these T4-NPs, mimicking the spikes of HIV-1 virion, exhibited exquisite targeting specificity to CD4+ cells and were efficiently taken up by the HIV-1 latency model J-Lat T cells, or the primary CD4+ T cells isolated from human peripheral blood mononuclear cells (PBMCs). This led to reactivation of proviral HIV-genome expression, viral protein production and virion release by the J-Lat T cells, and changes in primary T cells consistent with T cell activation. Intriguingly, the T4-NPs-induced activation, unlike the conventional shock-and-kill agents, was not global and did not involve the classic PKC (Protein Kinase C) or NFAT (Nuclear Factor of Activated T cells) pathways or production of cytokines typically associated with cytokine storm or other excessive immune responses. This new targeted, phage-based, approach could, in the future, be exploited to develop effective HIV-1 cure and other T-cell targeted therapies.

## Results

### Assembly of CD4-targeted CD4-DARPin-T4 Nanoparticles

In people living with HIV-1, activation of latent CD4+ T cells occurs when HIV-1 virions interact with the cell surface CD4 receptor through the envelope glycoprotein(Klasse and Moore 2004). We hypothesized that T4-NPs engineered to mimic such an interaction might also lead to activation of latent CD4+ T cells. To test this hypothesis, CD4-binding T4-NPs were prepared by decorating the capsid with a recombinant DARPin molecule that has high specificity and nanomolar binding affinity to human CD4 receptor(Schweizer et al. 2008).

Empty T4 capsid shells were prepared from *E. coli* infected with a phage mutant lacking Hoc, Soc, neck, and tail (Figure 1C)(Zhang et al. 2011). The recombinant CD4-DARPin fused to the N-terminal tip of Hoc was expressed in *E. coli*, purified, and displayed on T4 capsid through *in vitro* assembly (Figure 1D and 1E, Figure S1A and S1B).

Two versions of CD4-DARPin-Hoc fusions were constructed; one with a 6-aa linker (LYKYSD) between CD4-DARPin and Hoc, another with a longer 12-aa flexible linker ((GGSGGS)_2_) (Figure 1D). Both were displayed at equivalent efficiency, up to 1̃55 copies per particle, reflecting full occupancy. Capsids lacking the displayed CD4-DARPin, or those displayed with HIV-TAT, a nonspecific 14-aa Arg/Lys-rich cell penetration peptide (CPP) fused to Hoc, were used as controls. The highly basic TAT was previously shown to facilitate nonspecific attachment and delivery of cargo molecules across the negatively charged cell membrane(Milletti 2012; Tao et al. 2013). All the recombinant proteins were purified and displayed on the heads by *in vitro* assembly (Figure 1E).

New assays were developed to assess the targeting capability of CD4-DARPin-T4-NPs. First, an ELISA-based binding assay was developed to determine the binding specificity of CD4-DARPin-Hoc fusion proteins. Ninety-six well plates were coated with the soluble CD4 receptor extracellular domain (aa 1-370). Strong binding was observed only when the wells were exposed to CD4-DARPin-Hoc fusion protein with either the 6-aa linker or the 12-aa linker, but not to control Hoc or TAT-Hoc proteins (Figure 1F).

Another cell-based binding assay was developed to assess the targeting specificity to whole cells, by coating plates with CD4-DARPin or other test variants and exposing them to human CD4+ T cells (A3.01), CD4- T cells (A2.01), or CD4-overexpressing TZM-bl cells. After the unbound cells were washed off, the bound cells remaining in the wells were quantified using a sensitive CellTiterGlo luciferase assay that measured the amount of cellular ATP(Chand et al. 2017). Wells coated with CD4-DARPin-Hoc or the HIV-1 (JRFL strain) native-like SOSIP gp140 Env ectodomain trimers(AlSalmi et al. 2015; Ringe et al. 2013)efficiently bound to CD4+ A3.01 cells and the TZM-bl cells but not to CD4-lacking A2.01 cells (Figure 1G). The SOSIP gp140 trimers were purified, and their antigenicity was authenticated prior to binding(AlSalmi et al. 2015) (Figure S2). In contrast, the nonspecific TAT-Hoc showed approximately the same level of binding to both CD4+ and CD4- cells, whereas the negative Hoc-only control lacking CD4-DARPin showed no significant binding to any of the cells (Figure 1G). Additionally, externally added CD4-DARPin or CD4-specific monoclonal antibody (mAb) competitively blocked the binding of HIV-1 gp140 Env to CD4+ T cells (A3R5 cells), further demonstrating the binding specificity of the engineered proteins (Figure 1H).

Next, we tested if CD4-DARPin could target the T4-NPs to CD4+ HEK293T cells and deliver genetic payloads. The T4-NPs were first packaged with recombinant plasmid DNA containing expressible reporter genes *gfp* or *mCherry* using our *in vitro* DNA packaging system(Kondabagil, Zhang, and Rao 2006) and their surface was displayed with the targeting molecules, either the CD4-DARPin-Hoc or the gp140 Env trimers (Figure S2A-E) (Figure 1I-L). Both these T4-NPs when exposed to CD4+ cells allowed delivery and expression of the respective transgene in a CD4-targeted manner (Figure 1M and N). Furthermore, the GFP fluorescence was enhanced by the addition of chloroquine, suggesting that the NPs were internalized through an endocytosis-mediated process (Figure 1M). Chloroquine has been well-documented to acidify endosomes and enhance the escape of endosome-internalized payloads into the cytosol(Lonn et al. 2016). Additionally, when co-displayed with eGFP-Soc, the CD4-DARPin-T4-NPs but not the control NPs showed strong GFP fluorescence encircling the CD4+ cells at 60 minutes after exposure (Figure S3A-C).

The above sets of data demonstrate that we have functionalized the T4-NPs with targeting specificity to CD4+ human T cells that are permissive for HIV-1 infection.

### CD4-DARPin-T4-NPs activate J-Lat T cells and reverse HIV-1 provirus latency

We used J-Lat 10.6 full-length cells(Jordan, Bisgrove, and Verdin 2003), a well-characterized *in vitro* cell model of HIV-1 latency, to determine if the CD4-DARPin-T4-NPs can reactivate the resident HIV provirus. The J-Lat 10.6 cell line was originally developed by infecting immortalized Jurkat human T lymphocytes with a full-length HIV-1 vector that has a frame-shift mutation in the *env* gene and an insertion of *gfp* reporter gene in place of the *nef* gene. However, these cells are GFP fluorescence-negative because they are in a transcriptionally silent, post-integration latency state. If proviral reactivation occurred, they become GFP fluorescence-positive and produce noninfectious virus-like particles (VLPs) lacking the envelope(Jordan, Bisgrove, and Verdin 2003).

We prepared two types of CD4-targeted T4-NPs to test for reactivation; CD4-DARPin-T4-NPs and gp140 Env(trimers)-T4-NPs, as described above(Ananthaswamy et al. 2019; AlSalmi et al. 2015). We first established the targeting specificity of the ligands to J-Lat cells by our cell-binding assay described above (Figure S4A). Additionally, we demonstrated: i) loss of this binding upon down-regulation of CD4 expression by treatment with the latency reactivating agent PMA (Figure S4A), and ii) presence of red fluorescence around the cell when these the cells were exposed to T4-NPs co-displayed with mCherry protein (Figure S4B). These controls further attested the targeting specificity.

Remarkably, exposure of the CD4-DARPin-T4-NPs and Env(trimers)-T4-NPs to J-Lat cells resulted in abundant expression of GFP fluorescence in a dose-dependent manner (Figure 2). Maximum activation occurred at an m.o.i (ratio of NPs:cells) of 10^5^ (Figure 2A) and at 72 hours (Figure 2B), at which point most of the cells observed in the microscope appeared to be GFP-positive. Similar levels of activation were observed with Env(trimers)-T4-NPs, and also with the positive control, where the cells were treated with the shock-and-kill reagent PMA, which causes global activation of T cells (Figure 2A; see below).

**Figure 2.**
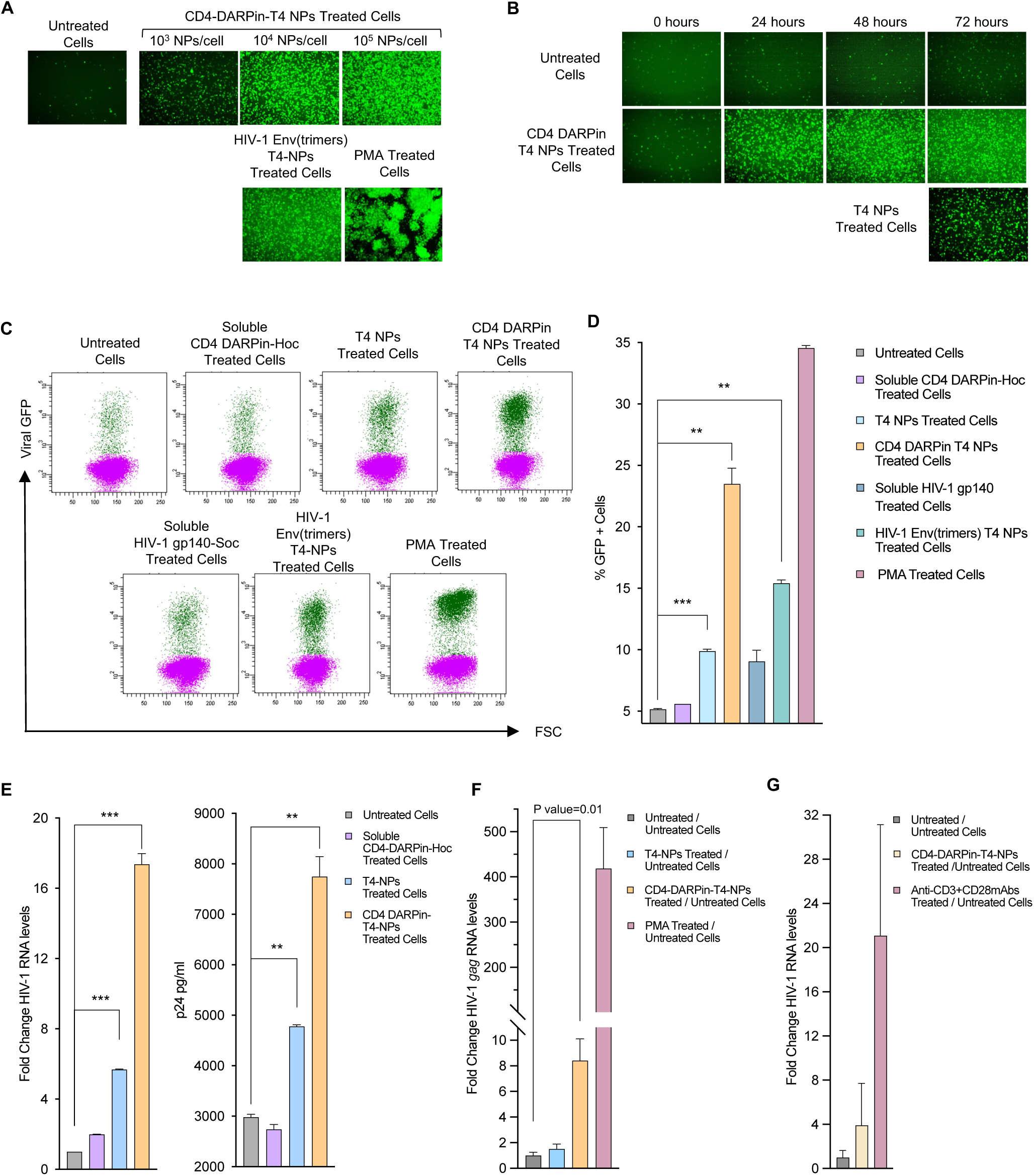
CD4-DARPin-T4-NPs activate latent J-Lat cells in a CD4-targeted manner. (A) GFP fluorescence micrographs showing the dose-dependent reactivation of latent cells by CD4-DARPin-T4-NPs, HIV-1 Env(trimers) T4-NPs, T4-NPs, and PMA. (B) GFP fluorescence micrographs showing the time-dependent reactivation of HIV-1 latent cells by CD4-DARPin-T4-NPs. (C) Flow cytometry dot plots showing HIV-1 reactivation profiles (GFP expression) in latent cells by CD4-DARPin-T4-NPs. Green and magenta dots represent GFP-positive and GFP-negative cells, respectively. The X-axis represents the frequency of GFP-positive cells, and the Y-axis represents cell size (FSC). (D) Quantification of GFP expression based on dot plots in panel C. Data are means ± S.D. of duplicate samples in two independent experiments. (E) Viral mRNA levels were quantified using real-time RT-PCR and normalized to beta-actin mRNA levels. The results were compared to unstimulated samples and shown as fold change (left graph). Data are means ± S.D of duplicate samples in two independent experiments. Gag-derived p24 HIV-1 protein in the cell culture supernatant assayed using a p24 ELISA (right graph). The average of three measurements ± S.D is shown. (F) HIV-1 genomic RNA levels in the cell culture supernatant assayed by real-time RT-PCR with RCAS RNA as an internal control. The results are represented as fold change from unstimulated samples. Data are means ± S.D. of duplicate samples and represent two independent experiments. The p-values were determined using a two-tailed, unpaired t-test. *** = P value <0.001 and ** = P value <0.01 and *= P value <0.05. (G) CD4-DARPin-T4-NPs target and activate resting CD4+ T cells isolated from ART-suppressed human donor PBMCs. Freshly isolated CD4+ CD25-resting T cells from ART-suppressed HIV-1 patients were left unstimulated or were stimulated with T4-NPs, CD4-DARPin-T4-NPs, anti-CD3/CD28 mAbs, or PMA. Each well was supplanted with 40 IU/ml of IL-2. After 10 days of incubation, the supernatants were analyzed for levels of genomic RNA and quantified by real-time RT-PCR, using RCAS RNA as an internal control. The results were compared to unstimulated samples and represented as fold change. Data are means ± S.D.

Flow cytometry confirmed these results (Figure 2C). Substantial levels of HIV proviral reactivation (green fluorescence) was observed with the targeted CD4-DARPin-T4-NPs (23.5%) and Env(trimers)-T4-NPs (15%), relative to 3̃4% with the PMA positive control. On the other hand, soluble CD4-DARPin or Env(trimers) (not displayed on T4) showed much lower stimulation (<10%), whereas the untreated and soluble Hoc controls were negative (Figure 2D).

An unexpected finding, however, was that the control T4-NPs lacking targeting molecules showed low but significant reactivation both by microscopy (Figure 2A and B) and by flow cytometry (1̃2%) (Figure 2D; also see below). This might be because the surface of T4 capsid exposing repetitive and symmetric structural elements mimics PAMPs (*p*athogen *a*ssociated *m*olecular *p*atterns) present on human viral pathogens. These might be recognized by the Toll-like receptors (TLRs) and activate the T cells. Additionally, small amounts of CpG phage DNA associated with the capsids(Zhang et al. 2011), a TLR9 agonist, might also have contributed(Iwasaki and Medzhitov 2004).

Together, the above results demonstrated that symmetric arrays of targeting ligands on the T4 capsid nanoparticle lattice can reactivate latent HIV-1 provirus present in J-Lat cells. Neither the soluble ligands nor the T4-NPs lacking the targeting ligand were effective.

### J-Lat cell activation by CD4-DARPin-T4-NPs leads to expression of proviral HIV-1 genes

Next, we investigated if the provirus reactivation as determined by GFP fluorescence also resulted in trascription of HIV-1 genes. HIV-1 transcripts were quantified by qPCR using viral-specific primers and correlated with GFP reporter expression. When compared to control unstimulated cells, the viral transcription increased by 17.4-fold when exposed to CD4-DARPin-T4-NPs vs 5.7-fold when exposed to T4-NPs lacking the targeting ligand and 1.99-fold with soluble CD4-DARPin-Hoc protein (Figure 2E). Additionally, a 2.6-fold increase in HIV gag/p24 antigen in the culture supernatant was observed with CD4-DARPin-T4-NPs versus 1.6-fold with T4-NPs, when compared to unstimulated cells (Figure 2E). The low-level stimulation observed with the control T4-NPS is consistent with the low-level reactivation/GFP fluorescence observed as above.

To determine if this was due to the secretion of assembled virus-like gag/p24 particles which should also contain packaged RNA genomes, the copy number of HIV-1 genomic RNAs in the culture supernatant was determined by multiplex qPCR using a primer-probe set for HIV-1 *gag* and RCAS(Replication Competent ALV LTR with a Splice acceptor) virus *gag,* (internal control). An 8-fold increase in the HIV-1 RNA levels was observed in the culture supernatant of cells treated with CD4- DARPin-T4-NPs when compared to the unstimulated cells (Figure 2F). Similar trends were observed with ART-suppressed patient-derived CD4+ CD25− resting T cells (Figure 2G).

These datasets show that the CD4-targeted T4-NPs reverse HIV-1 latency in J-Lat cells, leading to the expression of proviral HIV-1 genes, gag/p24 VLP assembly and release.

### Reactivation of HIV-1 latency by CD4-DARPin-T4-NPs does not involve the classic PKC and NFAT pathways

Physiologically, CD4 assists in the communication of T cell receptors (TCRs) with antigen-presenting cells (APCs). It binds to MHC-II on the APCs mediating downstream signaling that ultimately leads to T cell activation. Protein kinase C is a major regulator of these TCR signaling pathways(Isakov and Altman 2002). Therefore, we asked whether CD4-DARPin-T4-NPs act through the PKC signaling pathway to activate latent HIV-1 in J-Lat cells. As a control, we used the pan-PKC inhibitor Gö6983, which, as expected, completely suppressed the PMA-induced reactivation of latent HIV-1, as well as reduced the anti-CD3 and anti-CD28 co-stimulatory responses by 5̃0% (Figure 3A). Similarly, in another control, the inhibitor also suppressed the co-stimulatory effect by a mixture of anti-CD3 and anti-CD28 mAbs. However, Gö6983 exhibited no significant suppression of latent HIV-1 reactivation mediated by the CD4-DARPin-T4-NPs (Figure 3A).

**Figure 3.**
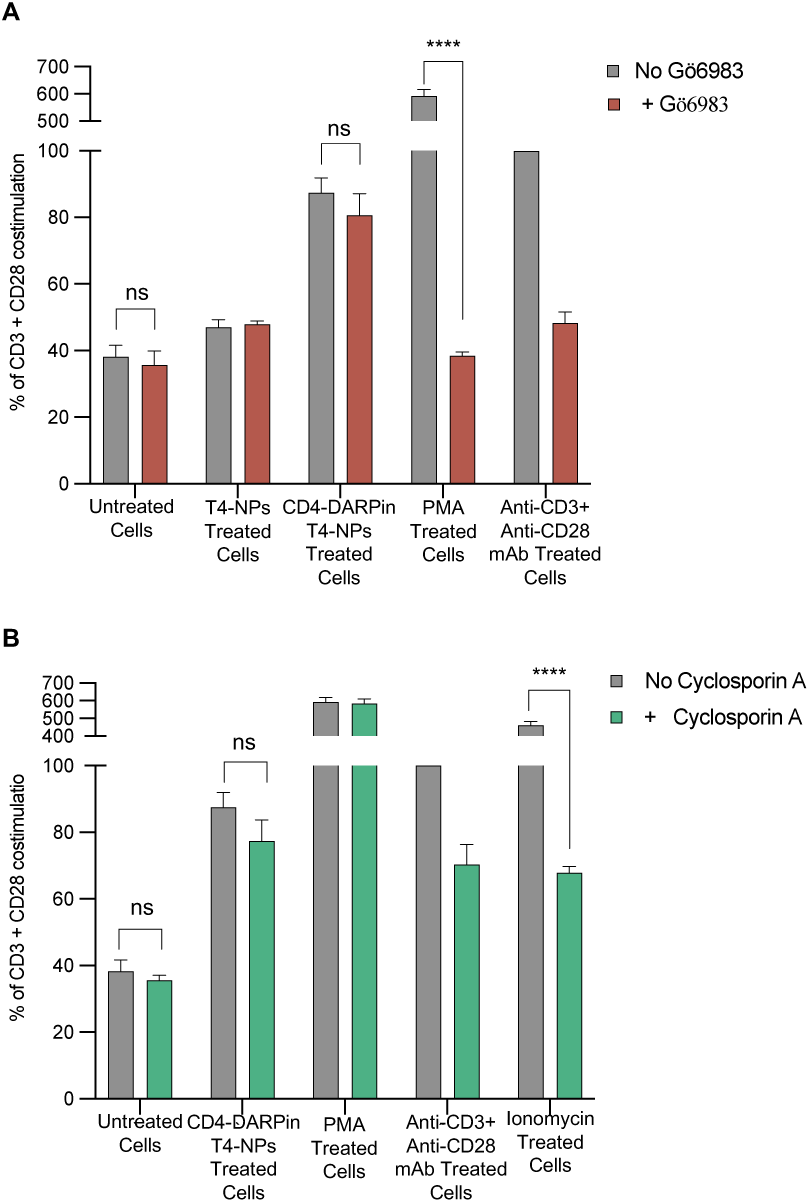
CD4-DARPin-T4-NPs reactivate latent HIV-1 independent of PKC and NFAT pathways. (A) Effect of a PKC inhibitor on the reactivation of latent HIV-1 by CD4-DARPin-T4-NPs. J-Lat 10.6 full-length cells were incubated with the PKC inhibitor Gö6983 for 1 hour prior to various treatments. Cells were collected and analyzed for GFP-positive cells after 48 hours of incubation. The results were normalized to the positive control, costimulation by anti-CD3 + anti-CD28 mAbs. Data are mean ±SD of duplicate samples. (B) Effect of cyclosporin A on the reactivation of latent HIV-1 by CD4-DARPin-T4-NPs. J-Lat 10.6 full-length cells were incubated with cyclosporin A 1 hour prior to each treatment. Cells were then collected and analyzed by flow cytometry for GFP-positive cells after 48 hours of incubation. Data are mean ± SD of duplicate samples. P-value was determined using a two-tailed, unpaired t-test. **** = P value <0.0001, ns; not significant.

NFAT, another essential transcription factor involved in T cell activation, is also reported to reactivate HIV-1 gene expression(Kinoshita et al. 1997). To assess whether NFAT played a role in the T4-NPs-mediated J-Lat cell activation, we used cyclosporin A (CsA), an inhibitor of the NFAT signaling pathway. We found that while CsA inhibited ionomycin-mediated activation of T cells though the NFAT pathway, no suppression of the CD4-DARPin-T4-NPs targeted reactivation of latent HIV-1 was observed (Figure 3B).

Finally, the reactivation by CD4-DARPin-T4-NPs did not induce cytokine storm as evident by no induction of Th1 and Th2 cytokines, or increased susceptibility to HIV-1 infection as evident by no stimulation of CCR5 co-receptor expression (Figure S6A and B).

Therefore, reactivation of HIV-1 latency in J-Lat cells is not a global activation, nor does it involve the classic PKC or NFAT pathways.

### CD4-DARPin-T4-NPs target T cells in human PBMCs and activate resting CD4+ T cells

Next, we have determined the targeting specificity of CD4-DARPin-T4-NPs to primary CD4+ T cells present in PBMCs of human donors. The human PBMCs were incubated with either the CD4-DARPin-T4-NPs or the PMA, and then stained for the T cell lineage and activation markers; CD3, CD4, CD38, and CD25 (Figure S5A).

Remarkably, this resulted in 98.8% depletion of CD4 staining on T cells, demonstrating exquisite targeting specificity of the CD4-DARPin-T4-NPs to primary human CD4+ T cells (Figure 4A). Furthermore, the treated T cells could no longer be stained with an anti-CD4 mAb because, presumably, the CD4 molecules now bound to CD4-DARPin-T4-NPs could no longer bind to the mAb. On the other hand, untreated cells or the cells treated with control T4-NPs laking the CD4-DARPin targeting molecule retained showed efficient binding by the anti-CD4 mAb. Additionally, both the CD4-DARPin-T4-NPs and the T4-NPs significantly induced the surface expression of the early activation marker CD38 on T cells (Figure 4B and FigureS5B and C), which was less evident for the late activation marker CD25 (Figure 4C and FigureS5D and E).

**Figure 4.**
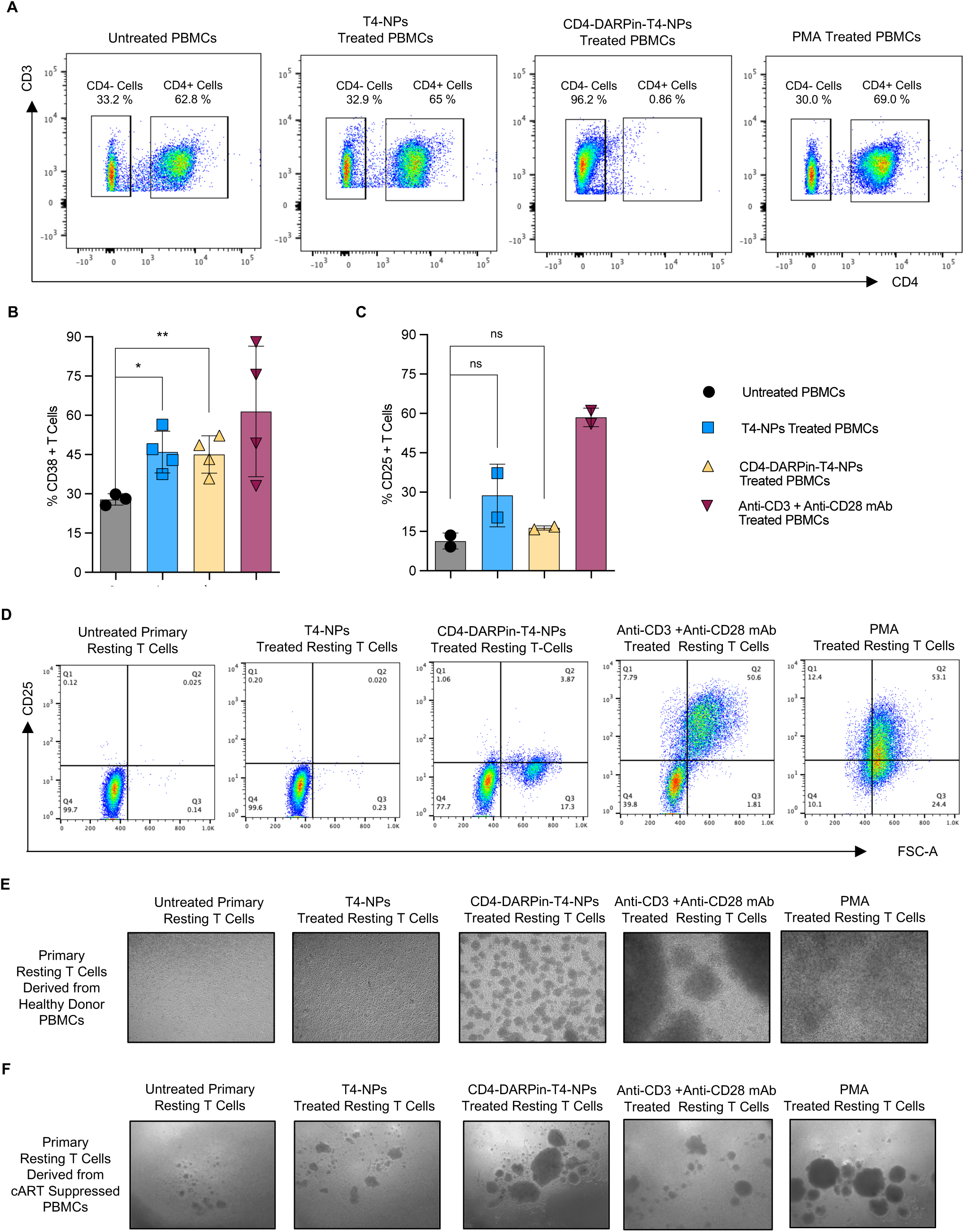
Displaying CD4-DARPin or HIV-1 Env trimers on T4 capsids targets the nanoparticles to CD4+ T cells. (A) PBMCs from healthy human donors were exposed to T4-NPs, CD4-DARPin-T4-NPs, and anti-CD3 + CD28 mAbs. After 48 hours of incubation, cells were co-stained for CD3, CD4, CD38, and CD25 receptors and analyzed by flow cytometry. (B and C) CD38(B) CD25(C) surface expression in CD3+ T cells. CD4+ CD25− resting T cells were isolated by negative selection from PBMCs of healthy human donors and exposed to T4-NPs, CD4-DARPin-T4-NPs, or anti-CD3 + anti-CD28 mAbs. After 48 hours of incubation, cells were analyzed for CD25 surface marker expression by flow cytometry. Two-tailed unpaired t-test, **=P value< 0.005, *=P value< 0.05. (D) CD25 surface expression and change in size c in the dot plots. Data are mean ± SD. (E) Clustering of resting CD4+ T cells isolated from healthy human donors PBMCs, following activation by targeted CD4-DARPin-T4-NPs. Freshly isolated CD4+ CD25− resting T cells were exposed to T4-NPs, CD4-DARPin-T4-NPs, anti-CD3/CD28 mAbs, or PMA. (F) CD4-DARPin-T4-NPs target and activate resting CD4+ T cells isolated from ART-suppressed human donor PBMCs. Freshly isolated CD4+ CD25− resting T cells from ART-suppressed HIV-1 patients were left unstimulated or were stimulated with T4-NPs, CD4-DARPin-T4-NPs, anti-CD3/CD28 mAbs, or PMA. Each well was supplanted with 40 IU/ml of IL-2. After 10 days of incubation, Microscopic images show cell clustering as an activation phenotype.

Since the latent HIV-1 reservoir is represented by the resting CD4+ T cells after infection, we assessed whether the CD4-DARPin-T4-NPs could target and activate this therapeutically more relevant fraction of T cells. Thus, we purified resting T cells (CD4+ CD25−) from PBMCs of a healthy human donor to 9̃8% purity by negative selection (Figure S6). These were either left unstimulated or stimulated with CD4-DARPin-T4-NPs or T4-NPs, while robust stimulation with CD3/CD28 mAbs served as a positive control. After 48 hours of incubation, a characteristic clustering of cells was observed in cells treated with CD4-DARPin-T4-NPs, similar to that observed in the positive control treated with CD3/CD28 mAbs (Figure 4D). We have also observed a weak upregulation of CD25 expression on the surface in 5̃% of the T cells that were treated with CD4-DARPin-T4-NPs, while no CD25 expression was detected in T4-NPs treated and untreated cells (Figure 4E). Additionally, about 20% of the cells treated with CD4-DARPin-T4-NPs exhibited an increase in size, as evident from the dot plots, similar to that observed in CD3/CD28 mAb-activated cells (Figure 4E). We have also observed a similar phenotype with the resting CD4+ T cells isolated from ART-suppressed patients when treated with CD4-DARPin-T4-NPs (Figure 4F). This phenotype represents T cell activation(Smith-Garvin, Koretzky, and Jordan 2009) and such clustering and increase in cell size have also been previously reported when CD4+ T cells were treated with the gp120 envelope protein that specifically binds to human CD4 receptor with high affinity(Kinet et al. 2002). No apparent morphological changes have been observed in the untreated cells, or cells treated with control T4-NPs lacking the CD4-DARPin targeting molecule (Figure 4E, F).

The above data demonstrated targeted activation of resting CD4+ T cells that constitute the T cell reservoir by the HIV-mimicking CD4-DARPin-T4-NPs.

## Discussion

In this paper, we describe a novel bacteriophage T4-based HIV mimic approach to target latent T cells with resident HIV-1 proviral genome and cause latency reversal and proviral reactivation. This is an essential step towards eliminating the latent viral reservoir in people living with HIV-1.

We set out by creating a HIV-1 envelope mimic by displaying a high-affinity human CD4 receptor binding ligand, CD4-DARPin, fused to the tip of a 18 nm-long flexible Hoc fiber. Up to 1̃55 such fibers decorated each T4 capsid nanoparticle. A series of datasets using ELISA, cell-binding, and flow cytometry assays demonstrated that these CD4-DARPin-T4-NPs bind to CD4 receptor-containing cells with exquisite specificity. This was evident not only with cultured CD4+ J-Lat 10.6 T cells, an *in vitro* model for HIV-1 latency, but also with primary T cells present in PBMCs from human donors. Remarkably, this resulted in near complete capture of CD4 receptors present on T cells because these cells could no longer be detectable by a CD4-specific mAb. When co-displayed with eGFP or mCherry, the bound NPs form a bright fluorescent layer around each CD4+ cell. Furthermore, they then delivered genetic payloads packaged inside the NPs into the host cell leading to efficient expression of the luciferase transgene.

Importantly, the CD4-DARPin-T4-NPs, or the T4-NPs decorated with the HIV-1 gp140 ectodomain trimers, caused robust proviral reactivation of J-Lat cells as determined by a series of assays demonstrating proviral GFP reporter expression, increased proviral transcription, and gag/gp24/viral RNA release into the culture supernatant. These results are consistent with the previously reported findings which implicated activation of CD4+ T cells by calcium-mediated signaling upon binding with the HIV-1 gp120 envelope protein, but not requiring viral entry(Perez et al. 1991).

The CD4 receptor plays a critical role in enhancing the TCR-mediated activation by interacting with MHC class II molecules on APCs. The cytoplasmic domain through interaction with the tyrosine kinase p56lck of the src family is known to initiate TCR signaling. Apparently, this CD4 function is activated by the observed interaction with CD4-DARPin or HIV-1 gp140. The CD4-DARPin-T4-NPs also showed activation of both CD3+ T cells and CD3− non-T cells which include adaptive T cells and innate cell types, respectively. This is evident from an increase in the surface expression of CD38 activation marker. Presumably, T4-NPs activated PBMCs through innate cell types such as dendritic cells and macrophages but this requires further investigation.

The activation by CD4-DARPin-T4-NPs was also recapitulated in primary resting CD4+ T cells isolated from human PBMCs, as shown by their clustering and increase in size upon exposure to the nanoparticles. Such a phenotype has been well-documented to be associated with T cell activation(Vasiliver-Shamis et al. 2009).

The observed non-global activation of J-Lat T cells that is independent of the classic PKC or NFAT pathways is intriguing and in clear contrast to the shock-and-kill reagents that cause massive global activation, even anaphylactic shock. It is also clear that activation by CD4-DARPin-T4-NPs does not induce cytokine storm or CCR5 co-receptor expression, also consistent with a non-global activation pathway. PKC was documented to be a master regulator of multiple signaling pathways such as AP-1, NF-*κ*B and ERK1/2. It seems that CD4-DARPin-T4-NPs bypass these pathways and use an alternative path that does not lead to full-blown global T cell activation. Further studies are needed to elucidate this potentially novel activation mechanism.

Our studies demonstrate that latency reversal and proviral reactivation required both CD4 receptor targeting and multipoint attachment to cell surface. Neither targeting alone using soluble CD4-DARPin or gp140, nor attachment without the targeting ligand, was found to be sufficient for maximal activation. This is reminiscent of the “viral plexes” formed through multipoint interactions between the virion’s Env spikes and the CD4 receptors on host cell surface, which is thought to be essential for membrane fusion and viral entry(Chen 2019). Our studies implicate that plexes formation with no associated viral entry might be sufficient, and is critically important, for signaling and activation that follows.

In conclusion, we show that engineered phage nanoparticles decorated with targeting ligands can serve as HIV-1 viral mimics by exhibiting remarkable targeting specificity to CD4+ T cells. At the same time, they can deliver and express transgenes, cause T cell activation that is independent of the detrimental global activation and/or cytokine storm, and reactivate the resident proviral HIV-1 gene expression and assembly and release of virus-like particles. Selective activation of T cells by phage T4-NPs nanoparticles, therefore, holds promise for further exploration and development of an effective HIV-1 cure. Since phage nanostructures are abundant in Nature and are highly engineerable(Zhu et al. 2023; Lemire, Yehl, and Lu 2018), it might be possible to design novel phage-based HIV-1 cure and T-cell targeted immunotherapies in the future.

## Methods and Materials

### Study participants

Patient-derived PBMCs was collected from HIV-infected individuals who were enrolled in a study of the effects of antiretroviral therapy in HIV-infected adults (clinical protocol 97-I-0082) at the National Institutes of Health (NIH) Clinical Center in Bethesda, MD.

### Purification of *10am-13am-hoc del*-soc del heads

The *10am-13am-hoc del-soc del* T4 phage mutant was constructed by standard genetic crosses and mutant heads were purified according to previously described protocols(Zhang et al. 2011). Briefly, *E. coli* P301 (sup-) cells (500 mL) infected with this mutant were lysed in 40 mL of Phosphate-Mg buffer (26 mM Na_2_HPO_4_/68 mM NaCl/22 mM KH_2_PO_4_/1 mM MgSO_4_, pH 7.5) containing 10 *μ*g/mL DNase I and chloroform (1 mL) and incubated at 37 °C for 30 min. The lysate was subjected to two low-speed (6,000 × g for 10 min) and high-speed (35,000 × g for 45 min) centrifugations, and the final pellet was resuspended in 200 *μ*L of Tris·Mg buffer (10 mM Tris·HCl, pH 7.5/50 mM NaCl/5 mM MgCl_2_) and purified by CsCl density gradient centrifugation. The major head band at about 1/3 from the top of a 5-mL gradient was extracted and dialyzed overnight against Tris·Mg buffer. The heads were further purified by DEAE ion-exchange column chromatography.

### Cell lines and reagents

J-Lat full-length clone 10.6, ACH-2, A3.01, A2.01, and TZM-bl cells were obtained through NIH AIDS Research and Reference Reagent Program(Jordan, Bisgrove, and Verdin 2003). Human PBMCs were obtained from StemCell Technologies Inc., CA (Catalog # 70025). Phorbol 12-myristate 13-acetate, PMA (16561-29-8), PKC inhibitor, Gö6983, (133053-19-7), Cyclosporin A, CsA, (C3362), and Ionomycin, (407952) were purchased from Sigma-Aldrich. Monoclonal anti-CD3 (catalog no. 555336) and anti-CD28 (catalog no. 555725) antibodies were obtained from BD Biosciences. As described, cells were cultivated as indicated in RPMI media (without glutamine, Invitrogen, 21870-076), Opti-MEM (Gibco, 31985-062) or DMEM; except where indicated, media were supplemented with Penicillin-Streptomycin-Glutamine (cat # 10378-016 Gibco-Thermo Fisher Scientific, MA) and 10% Fetal Bovine Serum (FBS, Quality Biologicals (Cat# 110-001-101HI). Reagents for nucleic acid isolation and qPCR from ART-suppressed patient-derived cells or culture supernatants; Glycogen, 1M Trizma hydrochloride buffer (Tris-HCl), 100% Isopropanol, Tris-buffered saline (TBS) tablets, recombinant DNaseI, 8M Guanidinium Hydrochloride (GuHCl), and 6M Guanidinium thiocyanate (GuSCN) were from Sigma-Aldrich (St. Louis, MO). Lightcycler 480 qPCR mix from Roche (Switzerland) was used. The Ribonuclease inhibitor was from Promega (Madison, WI). Superscript III Reverse Transcriptase, Deoxyribose nucleotide triphosphate mix (dNTPs), and Proteinase K were from Thermo Fisher Scientific (Waltham, MA). Oligos were purchased from Integrated DNA Technologies (IDT) (Coralville, IA).

### Plasmid construction

The plasmids pET-28b-Hoc and pET-28b-CPP-T-Hoc were constructed as previously described(Tao et al. 2013; Sathaliyawala et al. 2010). pET-28b-CD4-DARPin (55.2) was constructed by fusing 6X-Histidine tag to the N-terminus of CD4-DARPin 55.2 sequence from EMBL Nucleotide Sequence Database. The sequence was synthesized from Invitrogen and amplified using following primers FW1: 5’ATATACCATGGGCAGCAGCCATCATCATCATC3’ and BW1 5’TTAGGCTCGAGATTAAGCTTTTGCAGGATTTC having *NcoI* and *XhoI* cloning sites. The resulting fragment was purified by agarose gel electrophoresis, digested with appropriate restriction enzymes, and ligated with the gel-purified pET-28b vector DNA digested with the same restriction enzymes. For constructing pET28b-CD4-DARPin-LYKYSD-Hoc, two rounds of PCR were done. First round for the amplification of CD4-DARPin from pET-28b-CD4-DARPin-55.2 using primers, FW1 and BW2: 5’ GTCATATCGCTATATTTATACAGATTAAGCTTTTGCAGG 3’ and Hoc from the pET-28b-Hoc plasmid using FW2 5’ TAGGCTCGAGTTATGGATAGGTATAGATGAT-ACCAG TTTC 3’ and BW3: 5’CCTGCAAAAGCTTAATCTGTATAAATATAGCGATATGACTTTTA CAGTTGATATAACTC-CTAAAACACC3’. CD4-DARPin (GGGS)_2_GGSA-Hoc was amplified in similar fashion with primers having nucleotide sequence for (GGGS)_2_GGSA were used. Full-length CD4-DARPin-Hoc was acquired by the second round of PCR using the FW1 and BW3 primers and then digested with *NcoI* and XhoI. The digested fragment was subcloned into the pET-28b vector. Plasmids pAAV-GFP and pAAV-mCherry were purchased from Cell Biolabs. All the constructed plasmids were sequenced to confirm correct fragment insertion and accurate nucleotide sequence (Retrogen, CA).

### CD4-DARPin-Hoc protein expression and purification

Recombinant proteins produced in E. coli BL21 (DE3) RIPL cells were purified as per established protocols(Tao et al. 2013). Briefly, the BL21 (DE3) RIPL cells containing the recombinant plasmids were induced at 25 °C for 2 hours using 1 mM IPTG. The cells were collected via centrifugation (6,000 × g for 15 min at 4 °C) and then resuspended in 40 mL of HisTrap binding buffer, which contains 50 mM Tris·HCl at pH 8.0, 20 mM imidazole, and 300 mM NaCl. The cells were lysed using French-press (Aminco) and the soluble fraction containing the His-tagged fusion protein was isolated by centrifugation at 34,000 × g for 35 minutes. The supernatant was loaded onto a HisTrap column (GE Healthcare) and washed with 50 mM imidazole-containing buffer, and the protein was eluted with 20 to 500 mM linear imidazole gradient; The effluent was monitored at 280 nm and peak fractions The supernatant was loaded onto a HisTrap column (GE Healthcare) and washed with a buffer containing 50 mM imidazole. The protein was eluted using a gradient (linear) of imidazole from 20 to 500 mM. The effluent was monitored at 280 nm, and peak fractions were collected and purified by size exclusion chromatography, using a Hi-Load 16/60 Superdex-200 (prep-grade) gel filtration column (GE Healthcare). This was done in a buffer containing 25 mM Tris·HCl (pH 8.0) and 100 mM NaCl. The fractions were flash frozen in liquid nitrogen and stored at −80 °C.

### ELISA

ELISA plates (Evergreen Scientific, CA) were coated with 0.1 µg of protein per well in coating buffer [0.05 M sodium carbonate–sodium bicarbonate (pH 9.6)] overnight at 4°C. After washing three times with PBS-T buffer, the plates were blocked with PBS–3% BSA buffer for 1 hour at 37°C. The recombinant soluble CD4 (NIH AIDS Reagent program, 4615) was added and incubated for 1 hour at room temperature. The bound fraction was detected using anti-CD4 antibody Sim.2. (NIH AIDS Reagent Program, catalog # 723) by ELISA using soluble CD4 diluted 1:1000.

For P24 ELISA, Cell culture supernatant fluids were assayed for the Gag-derived p24 HIV protein using p24 ELISA kits (Zaptometrix) and by following manufacturer’s instructions.

### Cell binding assay

The cell binding assay was performed as described before(Chand et al. 2017); briefly, purified CD4-DARPin variants and gp140 Env proteins were coated onto a 96-well black, clear-bottom plate (Greiner), 15 poles per well, for 1 hour. The wells were subsequently washed 3 times with blocking buffer (1mM MnCl_2_, 0.1mM CaCl_2_, 10mM HEPES, 150 mM NaCl, and 10% FBS) and then incubated with the blocking buffer for 1 hour. Wells were then washed 3 times with wash buffer (1mM MnCl_2_, 0.1mM CaCl_2_, 10mM HEPES, 150mM NaCl, and 1% FBS). A3.01 (CD4+), A2.01 (CD4−), TZM-bl and J-Lat 10.6 full length clone cells (50 *μ*l/well of 4×10^6^ cells per ml) were added in cell dilution buffer (wash buffer containing 5% FBS) and allowed to bind for 1 hour. Wells were then washed 5 times with wash buffer and the remaining bound cells were detected with the CellTiterGlo kit (Promega cat #G7570) as per the manufacturer’s instructions.

### In vitro DNA packaging and protein display on T4 heads

For in vitro DNA packaging assays(Rao, Thaker, and Black 1992), each 20 µl of reaction mixture contained purified T4 heads (about 2 × 10^10^ particles), purified full-length gp17 (3 µM), and linearized DNA in packaging buffer [30 mM Tris-HCl (pH 7.5), 100 mM NaCl, 3 mM MgCl_2_, and 1 mM adenosine 5’ triphosphate (ATP)]. The mixture was incubated at 37°C for 30 min, followed by benzonase nuclease addition and incubation at 37°C for 20 min to remove excess unpackaged DNA. The encapsidated nuclease-resistant DNA was released by treatment with 50 mM EDTA, proteinase K (0.5 g/µl; Thermo Fisher Scientific, MA), and 0.2% SDS for 30 min at 65°C. The packaged DNA was analyzed by 1% (w/v) agarose gel electrophoresis followed by staining with ethidium bromide, and the amount of packaged DNA was quantified using Quantity One software (Bio-Rad, CA). The packaging efficiency was defined as the number of DNA molecules packaged per T4.

After encapsidating linearized DNA as described above, T4 heads were incubated with Soc- and/or Hoc-fused proteins at 4°C for 45 min. The mixtures were sedimented by centrifugation at 30,000g for 45 min, and unbound proteins in the supernatants were removed. After washing twice with PBS, the pellets were incubated at 4°C overnight and then resuspended in PBS for SDS–polyacrylamide gel electrophoresis (SDS-PAGE) analysis or Opti-MEM for transduction. After Coomassie Blue R-250 (Bio-Rad, CA) staining and destaining, the protein bands on SDS-PAGE gels were scanned and quantified by laser densitometry (Personal Densitometer SI; GE Healthcare, IL). The band densities of the Hoc, Soc, and gp23 were determined for each lane separately, and the copy numbers of bound Hoc or Soc fusion molecules per capsid were calculated using the major capsid protein gp23* as the internal control (930 copies per capsid).

### Cell transduction and the detection of gene delivery

CD4+HEK293T cells were seeded in 24-well plates at 2 × 10^5^ cells per well in complete DMEM. After 24 hours, the cells were incubated with the T4, CD4-DARPin-T4 vectors at different m.o.i. in antibiotic-free Opti-MEM for 6 hours. Thereafter, Opti-MEM was removed and replaced with complete DMEM. The cells were further incubated at 37°C for an additional 48 hours. GFP trans-gene expression was observed by fluorescence microscopy (Carl Zeiss, Germany) at 48 hours after transduction. To analyze luciferase gene delivery into cells by T4 or CD4-DARPin-T4, we measured luciferase activity with the Luciferase Assay System (Promega, WI, Cat # E1500) according to the manufacturer’s instructions. Briefly, the growth medium was removed, and cells were rinsed with PBS buffer. After removing the wash buffer, 150 *μ*l of passive lysis buffer was added to each well, followed by gentle shaking at RT for 20 mins. 20 *μ*l of the cell lysate was then transferred to a 96-well white opaque plate and mixed with 80 *μ*l of Luciferase Assay Reagent, and the luminescence signal was recorded using the GloMax-Multi Detection System (Promega, WI).

### Cytokine secretion analysis

Fresh PBMCs (5 × 10^5^ cells per well) were left unstimulated or stimulated with T4-NPs, CD4− DARPin-T4-NPs, and anti-CD3 & anti-CD28 Abs for 48 hours at 37°C. Medium-treated PBMCs served as a negative control. After stimulation, cell-free supernatant was collected and analyzed by Bio-Plex Pro^TM^ Human Cytokine Th1/Th2 Assay according to the manufacturer’s instructions (Cat #M5000005L3 Bio-Rad). All tests were performed in duplicates, and mean values were calculated.

### Resting CD4+ T cell subset isolation

Primary human CD4+ T cells were isolated from PBMCs using a EasySep™ Human Resting CD4+ T Cell Isolation Kit, according to the manufacturer’s instructions (Cat #17962 STEMCELL Technologies). Briefly, cryopreserved PMBCs were thawed, washed, and resuspended at 5 × 10^7^ cells/mL in PBS containing 2% FBS. To the sample, EasySepTM Human Resting CD4+ T Cell Isolation Cocktail, CD25 Depletion Cocktail, and Rapid Spheres were added, followed by incubation for 5 mins at room temperature. After incubation, the sample volume was adjusted to 2.5 ml with buffer containing PBS with 2% FBS and 1mM EDTA, PBS should be without Ca2+ and Mg2+. The tube was placed into the magnet for 5 mins at RT and poured into a new tube to obtain the enriched cell suspension. Purified CD4+ T cells were >97% pure as assessed by flow cytometry.

### Measurement of the reactivation of latent HIV-1 in J-Lat cells

2.5 × 10^5^ J-Lat 10.6 full-length clone cells were resuspended in OPTI-MEM media in flat bottom 24 well plate. The cells were then treated with the indicated concentrations and the number of activators or phage T4-NPs, respectively. After 6 hours, FBS was added to a final concentration of 10%. Stimulation with 2.5 *μ*g/ml anti-CD3 and 1 *μ*g/ml anti-CD28 monoclonal antibodies were used as positive controls. After 40-48 hours at 37°C, reactivation of latent HIV-1 was determined by quantifying the percentage of GFP+ cells using the FACS ARIA and analyzed using FlowJo (BD Biosciences) software.

### HIV-specific mRNA measurements

RNA was isolated using Direct-Zol RNA MiniPrep Plus (Zymo Research, Irvine, CA) and cDNA was synthesized by using iScript cDNA Synthesis Kit (Bio-Rad Laboratories, Hercules, CA). qRT-PCR was performed on the Applied Biosystems StepOne™ Real-Time PCR System using iTaq™ Universal SYBR® Green Supermix (Bio-Rad Laboratories, Hercules, CA) with the HIV-1 mRNA primer set: - 5’GTGTGCCCGTCTGTTGTGTGA-3’ primer 2: 5-GCCACTGCTAGAGATTTTCCA-3 and the GAPDH primer set: - 5’AAGGTGAAGGTCGGAGTCAAC-3’ and - 5’GGGGTCATTGATGGCAACAATA-3’. The following cycling conditions were used for all qRT-PCR reactions: 95°C for 10 minutes, followed by 40 cycles at 95°C for 15 seconds and 60°C for 1 minute. Relative fold changes were calculated after normalization using GAPDH as a reference gene.

### Immunofluorescence assay

Cells were cultured on glass coverslips and fixed at room temperature using 3.7% PFA for 10 minutes. After fixation, cells were incubated for 1 hour with either CD4-DARPin-T4-NPs or T4-NPs. The coverslips were then mounted on slides with DAPI-conjugated mounting media. Labeled cells were observed via epifluorescence using an Olympus BX60 Fluorescence Microscope (OPELCO, Dulles, DC, USA) equipped with a UPlanFl 60×/NA 1.3, phase 1, oil immersion objective. Images were captured utilizing an HQ2 CoolSnap digital camera (Roper Scientific, Berlin, Germany) and Metamorph Imaging software (Molecular Devices, Sunnyvale, CA, USA). Image processing and compilation were conducted with Image J (National Institutes of Health).

### Transfections of HIV-1 Env-Soc Trimers

HEK293F suspension cells were grown overnight to a density of 10^6^ cells/ml. Large-scale transfection in 1L culture volume was done using FreeStyle MAX transfection reagent (Life Technologies). Briefly, the cells were transfected with 1 *μ*g of gp140-Soc plasmid DNA/ million cells. As these clones are cleavage-sensitive, the cells were co-transfected with the furin plasmid at a gp140-Soc: furin plasmid DNA ratio of 3:1 to ensure near 100% cleavage. Plasmid DNAs and MAX reagent were diluted in OptiPRO SFM medium, mixed and incubated for 10 min at room temperature. The mixture was then added to the HEK293F suspension cells. After 12 hours, the transfected cells were supplemented with 100 ml of fresh HyClone SFM4HEK293 medium (GE Healthcare) and sodium butyrate (Reeves et al., 2002) solution (Sigma-Aldrich; final concentration of 2 nM). On day 5, the supernatant was harvested by centrifuging the cells and filtered using a 0.2 *μ*m filter (Corning, Inc.) for purification of the secreted protein.

### HIV-1 gp140-soc protein purification

Secreted twin strep-tagged gp140-soc proteins in the harvested and filtered supernatant (1L) were supplemented with protease inhibitor tablets (Roche Diagnostics) to prevent protein degradation and 5 ml of BioLock biotin blocking solution (iba Life Sciences GmbH) to mask the biotin present in the supernatant. Next, the supernatant was loaded onto a 1 ml StrepTactin column (Qiagen) at a flow rate of 0.7 ml/min in the AKTA prime-plus liquid chromatography system (GE Healthcare). Non-specifically bound proteins were removed by passing at least 20 column volumes of wash buffer (50 mM Tris-HCl, pH 8, and 300 mM NaCl) until the 280nm absorbance reached the baseline level. Bound gp140-soc proteins were eluted with StrepTactin elution buffer (5 mM d-Desthiobiotin, 25 mM Tris-HCl, pH 8, with 150 mM NaCl) at a flow rate of 1 ml/min. Effluent was monitored at 280 nm and peak protein fractions were pooled and loaded onto the size-exclusion chromatography column using Hi-Load 16/60 Superdex-200 (prep-grade) gel filtration column (GE Healthcare) in a buffer containing 25 mM Tris·HCl (pH 8.0) and 100 mM NaCl. All the eluted fractions were collected and analyzed on BLUE Native PAGE and reducing/non-reducing SDS-PAGE for biochemical analysis.

### StrepTactin ELISAs

SEC-purified HIV-1 Env-Soc trimers was tested for antigenicity using ELISA involving Microplates pre-coated with StrepTactin (IBA Life Sciences GmbH). Microplates were coated with 1 *μ*g/ml SEC-purified gp140-Soc trimers in 100 *μ*l/well of coating buffer (25 mM Tris-HCl, pH 7.6, 2 mM EDTA, and 140 mM NaCl) for 2 hours at room temperature. After washing with PBST (0.05% Tween 20 in PBS), the trimers were incubated with a series of diluted PGT145 and 8ANC195 antibodies for 1 hour at 37°C. The antibodies were sourced from the NIH AIDS Research and Reference Reagent Program. Following incubation, unbound antibodies were washed away, and the bound antibodies were detected using HRP-conjugated rabbit anti-human antibodies (Santa Cruz Biotechnology). Peroxidase substrate (TMB microwell peroxidase substrate system, KPL) was added and reaction was terminated by BlueSTOP solution (KPL) and optical density at 650 nm was measured (Molecular Devices).

### Isolation of HIV RNA from cell culture supernatants

CD4+ T cells from ART-suppressed patients were either left untreated or treated with T4-NPs, CD4-DARPin-T4-NPs, anti-CD3 and anti-CD28 mAbs, or PMA. HIV RNA from free virions released into the supernatant was extracted every three days for a maximum of nine days. The cryopreserved supernatants were rapidly thawed at 37°C, and 500 *μ*l of each was diluted in Tris-buffered saline (TBS) before being centrifuged at 21,100 x g for one hour at 10°C to pellet the cell-free virus. After removing the supernatant, the pellet was resuspended in 100 ml of 3M guanidinium hydrochloride (GuHCl) with 50 mM Tris-HCl pH 7.6, 1 mM calcium chloride, and 100 mg proteinase K, and incubated at 42°C for one hour. Subsequently, 400 ml of 6 M guanidinium thiocyanate (GuSCN) (containing 50 mM Tris HCl pH 7.6, 1 mM EDTA, and 600 mg/mL glycogen) was added, followed by another incubation at 42°C for 10 minutes. After this, 500 ml of room temperature 100% isopropanol was incorporated into the guanidinium mixture, which was inverted 20 times and centrifuged at 21,000 x g for 15 minutes at room temperature to pellet the RNA. The supernatant was discarded, and the pellet washed with 70% ethanol. Finally, the RNA pellets were air-dried for five minutes and stored at −80°C for downstream assays.

### cDNA synthesis and RT-qPCR using patient samples

Complementary DNA (cDNA) was synthesized from 10 *μ*l of HIV RNA transcripts or extracted CA-RNA using random hexamers and SuperScript IV Reverse transcriptase according to manufacturer protocol (Thermo Fisher). Next, a multiplexed qPCR master mix was made with a final concentration of 1X Lightcycler 480 Probes Master Mix (Roche, Switzerland), 600 nM forward and reverse primers, and 100 nM probe. We used a primer-probe set HIV-1 gag and RCAS gag. The copy number was determined from the standard curve generated from HIV RNA transcript standards that were diluted from 10 x 10^6^ to 1 copy/per well and assayed in triplicates for each run.

HIV iSCA Forward Primer 5’ TTTGGAAAGGACCAGCAAA-3’

HIV iSCA Reverse Primer 5’ CCTGCCATCTGTTTTCCA-3’

HIV iSCA Probe 5’ 6FAM-AAAGGTGAAGGGGCAGTAGTAATACA-TAMRA-3’

RCAS Forward Primer 5’ GTCAATAGAGAGAGGGATGGACAAA-3’

RCAS Reverse Primer 5’ TCCACAAGTGTAGCAGAGCCC-3’

RCAS Probe 5’ FAM-TGGGTCGGGTGGTCGTGCC-TAMRA-3’

## Statistical Analyses

For the statistical comparison of two individual groups, a Student’s t-test was performed with two-tailed distributions and equal variance. For comparison of multiple groups, a two-way ANOVA test was performed using GraphPad Prism 6.05 Software (GraphPad Prism Software, Inc.).

## Supplementary Figures

**Figure S1.**
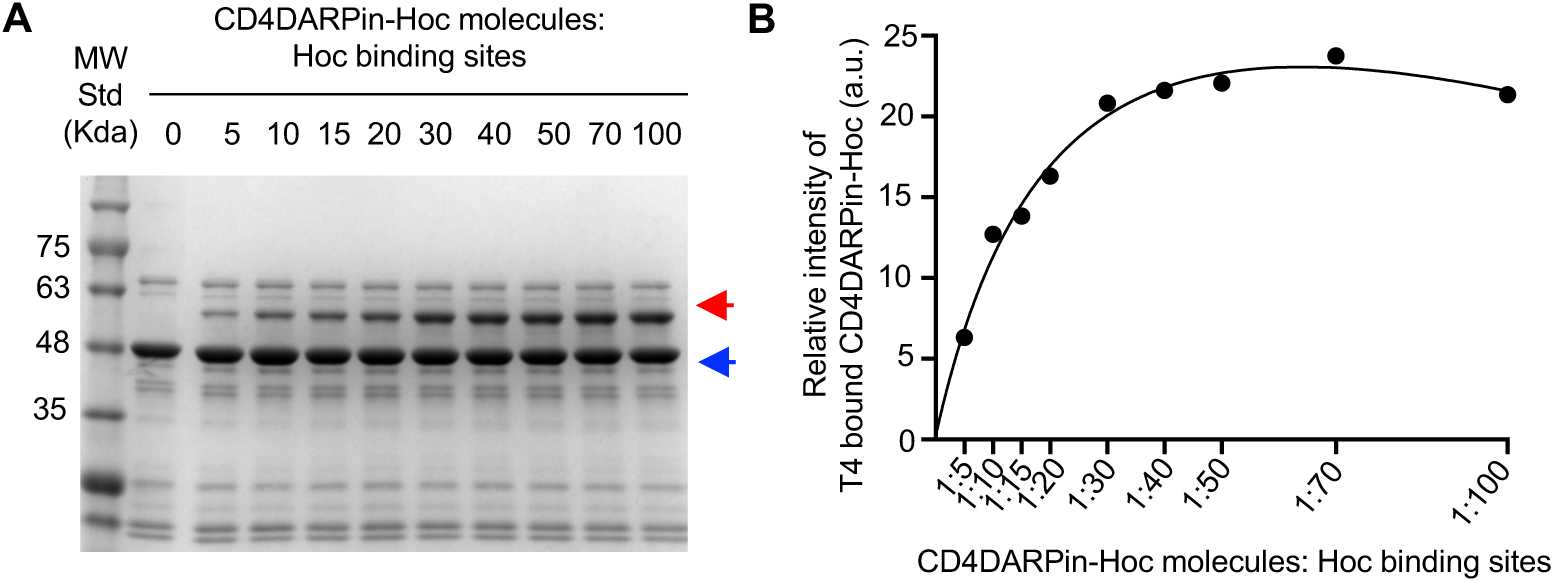
Assembly of CD4-DARPin-T4-NPs. (A) Display of CD4-DARPin-Hoc on the T4 capsid at increasing ratios of CD4-DARPin-Hoc molecules to Hoc binding sites (5:1 to 100:1). The red and blue arrows show the positions of bound CD4-DARPin-Hoc and major capsid protein gp23*, respectively. (B) CD4-DARPin-Hoc binding curve with nonlinear regression. The density volumes of T4-bound CD4-DARPin-Hoc were quantified by laser densitometry of the SDS-PAGE results in panel A. Similar results were obtained when the experiment was carried out with Hoc, CD4-DARPin-12a.a-Hoc, and Hoc-TAT proteins.

**Figure S2.**
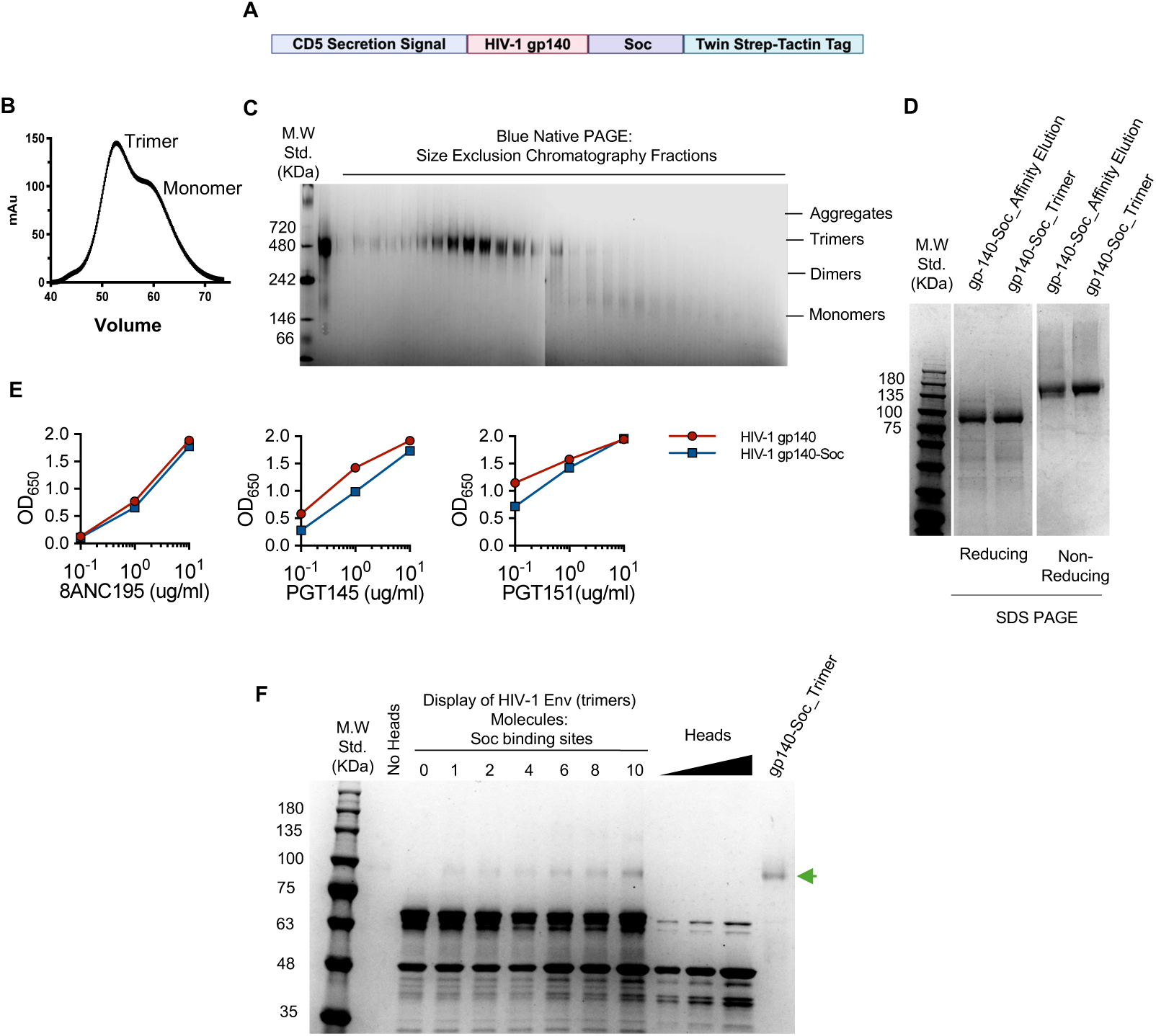
Assembly of gp140 HIV-1 Env trimers on T4 capsids. (A) Schematic of HIV-1 gp140-Soc fusion construct. The codon-optimized HIV-1 gp140 (JRFL) coding sequence was fused to the N-terminus of Soc via a linker. (B) Size exclusion chromatography elution profile. (C) Blue-NATIVE (non-denaturing) PAGE analysis of the eluted fractions. (D) Reducing and non-reducing SDS-PAGE of affinity chromatography-purified sample and size exclusion chromatography-purified gp140 Env trimers. (E) Antigenicity of gp140-soc Env trimers. Comparative binding of cleaved gp140 trimers and gp140-soc trimers is depicted with trimer-specific mAbs 8ANC195(Top) and PGT145 (Bottom)(AlSalmi et al. 2015; Ananthaswamy et al. 2019). Trimers were captured on the Strep-Tactin ELISA plates through the C-terminal StrepTag II at a constant concentration of 1 µg/ml for the assay. Each panel shows the binding curve from three replicates at the indicated concentrations of the mAbs. No statistically significant differences in binding were observed between the gp140-soc trimers and gp140 trimer pairs using an unpaired two-tailed t-test. (F) Display of HIV-1 gp140-Soc trimers on T4 heads. About 4 × 10^10^ Hoc-Soc-head particles were incubated with increasing ratios of HIV-1 gp140-Soc molecules to Soc-binding sites on the capsid (0:1–10:1 as indicated). After washing off the unbound protein, the samples were electrophoresed on a 4-20% (wt/vol) gradient SDS/PAGE. The position of the HIV-1 gp140-Soc protein is marked with a green arrow.

**Figure S3.**
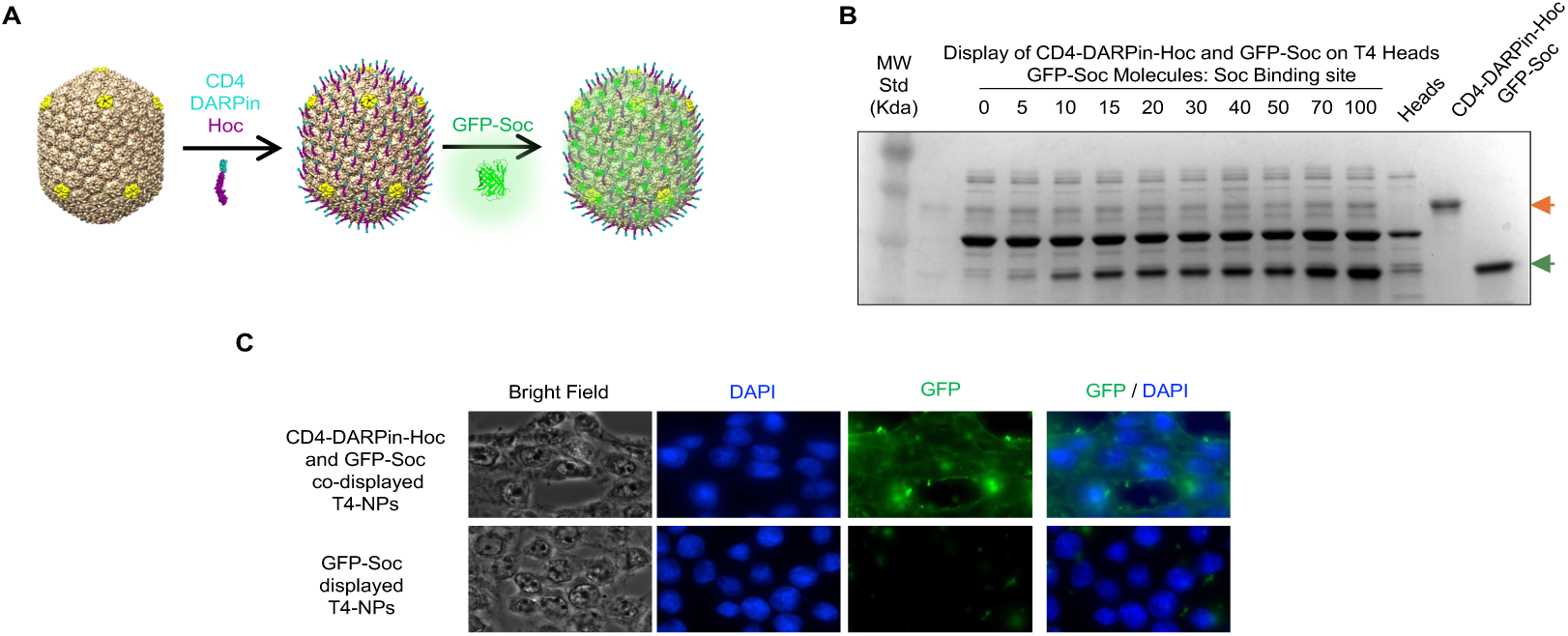
Targeting of CD4-DARPin-Hoc and GFP-Soc co-displayed T4 heads to CD4+ cells. (A) Schematic of CD4-DARPin-Hoc and eGFP-Soc co-display on T4 heads. (B) SDS-PAGE showing co-display. About 4 × 10^10^ Hoc-Soc-heads were incubated with a 30:1 ratio of CD4-DARPin-Hoc molecules: Hoc binding sites, and 0:1–100:1 ratio of mCherry-Soc molecules to Soc-binding sites. After washing off the unbound protein, the samples were electrophoresed on a 4-20% gradient SDS-PAGE. The positions of the CD4-DARPin-Hoc protein (orange) and mCherry-Soc (pink) are marked with arrows. (C) CD4+ HEK293T cells were grown on coverslips and fixed by paraformaldehyde, followed by incubating with either CD4-DARPin-Hoc displayed or CD4-DARPin-Hoc and eGFP co-displayed T4-NPs for 60 mins followed by washing and fixing with DAPI-mount media and fluorescence microscopy. Images are at 60X magnification.

**Figure S4.**
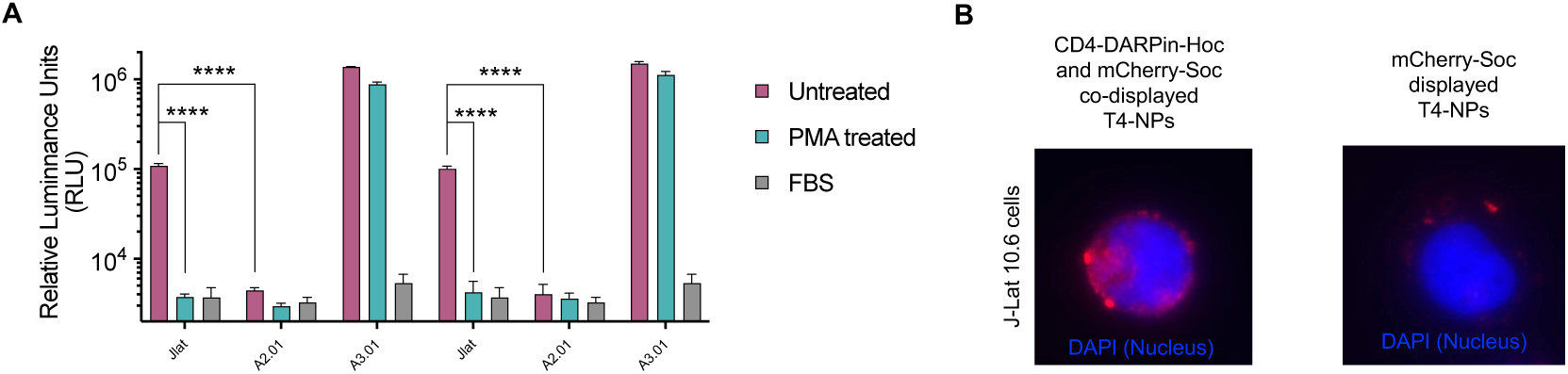
Targeting of CD4-DARPin-Hoc and mCherry-Soc co-displayed T4 heads to J-Lat cells. (A) CD4-DARPin-Hoc and HIV-1 gp140 Env trimers were coated onto a 96-well plate and blocked with FBS to prevent non-specific binding. The J-Lat 10.6 full-length cells were added to the wells and the extent of binding was quantified by the chemiluminescence-based assay (CellTiteGlo Kit, Promega). (B) J-Lat 10.6 full-length cells were grown on coverslips and fixed, followed by incubating either with CD4-DARPin-Hoc and mCherry-Soc co-displayed or CD4-DARPin-Hoc displayed T4-NPs for 60 mins followed by washing and fixing with DAPI-mount media and fluorescence microscopy. Images are at 60X magnification.

**Figure S5.**
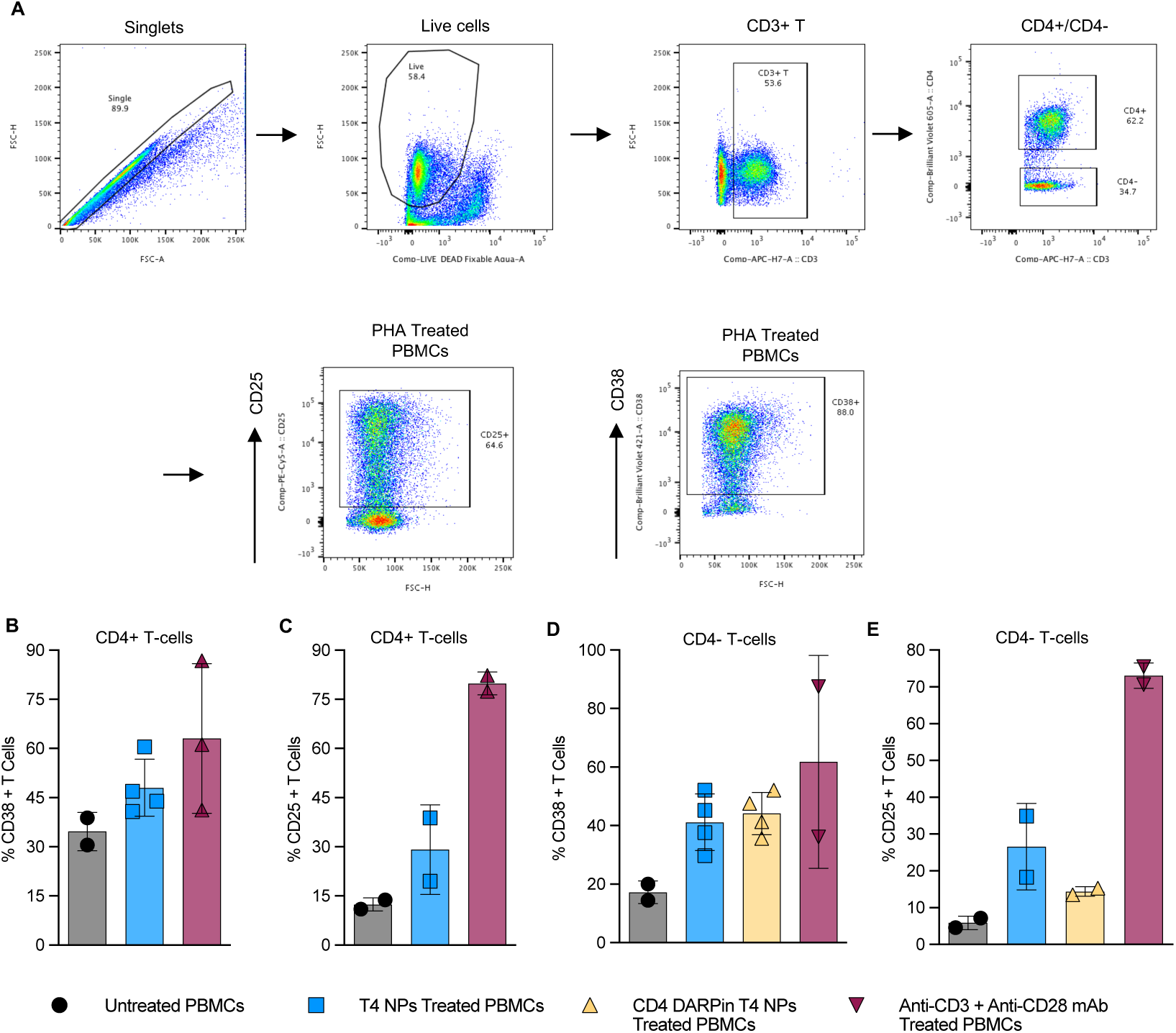
Activation of T cells in human PBMCs by CD4-DARPin-T4-NPs. PBMCs from healthy human donors were exposed to T4-NPs, CD4-DARPin-T4-NPs, or anti-CD3+ anti-CD28 mAbs. After 48 hours of incubation, cells were stained for viability dye, CD3, CD4, CD25, and CD38, followed by flow cytometry analysis, as shown in Figure 4. (A) Flow cytometry plots detailing gating strategy. (B and C) CD38 and CD25 surface expression on CD4+T cells. (D and E) CD38 and CD25 surface expression on CD4-T cells.

**Figure S6.**
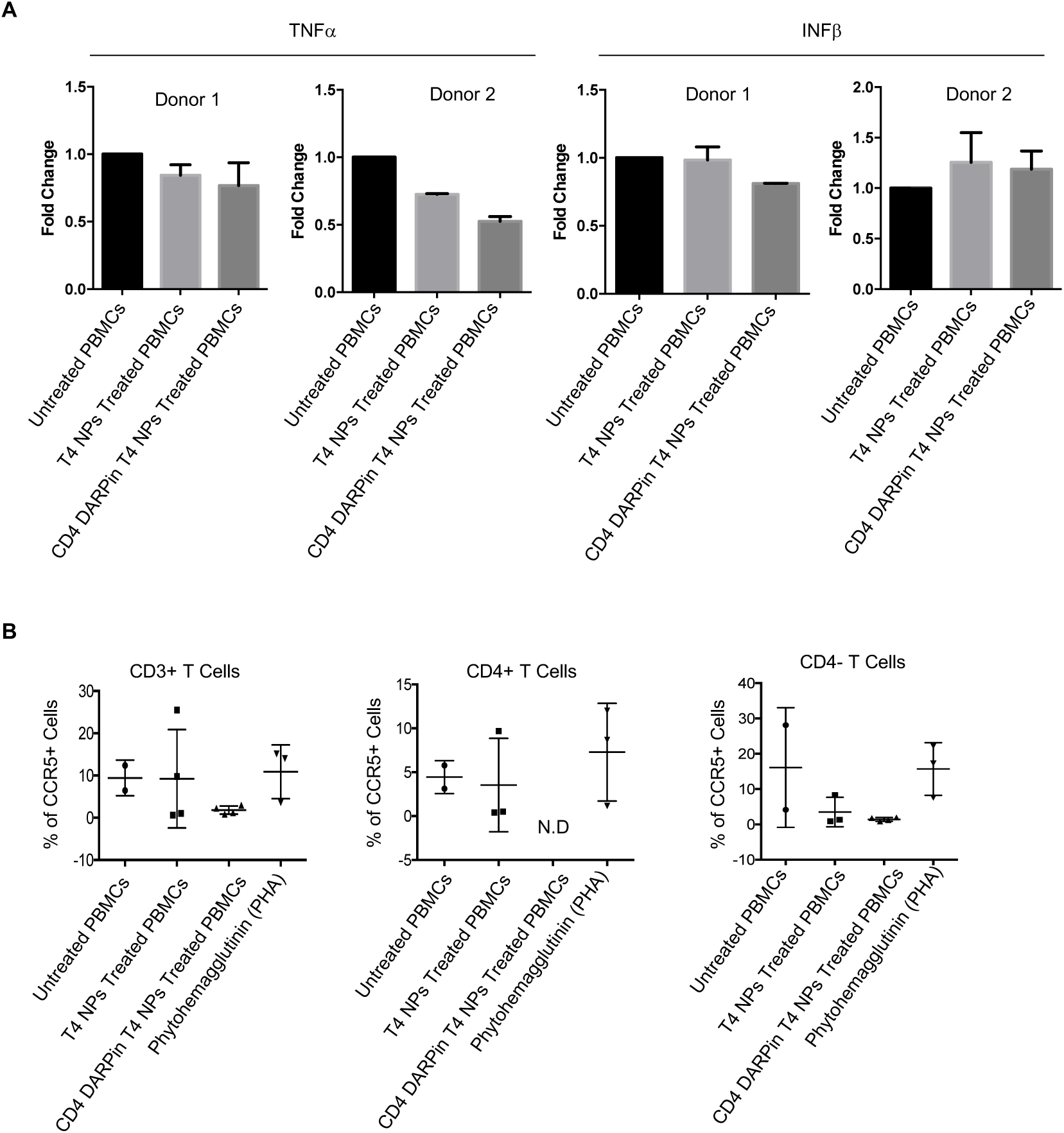
CD4-DARPin-T4-NPs do not induce cytokine storm or enhance CCR5 expression on T cells. Freshly isolated human PBMCs were exposed to T4-NPs, CD4-DARPin-T4-NPs and PHA for 48 hours. (A) Cells were analyzed for TNF-a and INF-b mRNA levels and quantified using real-time RT-PCR. The results were compared to unstimulated samples and showed as fold change. Data are means ± S.D, P value <=0.001, two-tailed, unpaired t-test. (B) Cells were analyzed for CCR5 expression on T cells by flow cytometry. No significant change was observed between the unstimulated and either type of T4-NPs. N.D. not detected. Data are shown as means ± S.D.

## Data Availability

Requests for further information and reagents should be directed to, and will be fulfilled by the lead contact, Venigalla Rao.

## Acknowledgments

We thank Masashi Waga and Victor Padilla-Sanchez [The Catholic University of America (CUA)] for assistance in preparing the T4 capsid images. This work was supported by BtB grant (to F.M. and V.B.R.), NIDA grant DP1DA060580-01 (to V.B.R.) and NIAID grant R01AI175340 (to V.B.R.) and NIAID grants R01AI145666 and R01AI157852 (to H.H).

## Author Contributions

V.R. directed the project. H.B. and V.R. conceived and designed the study. H.B., J.Z., S.J., N.A., M.M., P.T., C.L., S.H., and C.Z performed experiments. H.B. H.H., F.M., and V.R. analyzed and interpreted the data. V.R. and H.B. wrote the manuscript. F.M., H.H., and M.K. edited the manuscript.

## Decleration of Interests

V.R. and H.B. are co-inventors of a patent application (17/736,181) describing the engineered bacteriophage T4 nanoparticles for targeted therapies.

